# Distinct roles of two RDL GABA-receptors in fipronil action in the diamondback moth (*Plutella xylostella*)

**DOI:** 10.1101/2020.08.17.255026

**Authors:** Benjie Li, Kunkun Wang, Dongping Chen, Ying Yan, Xuling Cai, Huimin Chen, Ke Dong, Fei Lin, Hanhong Xu

## Abstract

The phenylpyrazole insecticide, fipronil, blocks insect RDL γ-aminobutyric acid (GABA) receptors, thereby impairs inhibitory neurotransmission. Some insect species, such as the diamondback moth (*Plutella xylostella*), possess more than one *Rdl* gene. The involvement of multiple *Rdls* in fipronil toxicity and resistance remain largely unknown. In this study, we investigated the roles of two *Rdl* genes, *PxRdl1* and *PxRdl2*, from *P. xylostella* in the action of fipronil. Expressed in *Xenopus* oocytes, *Px*RDL2 receptors were 40-times less sensitive to fipronil than *Px*RDL1. *Px*RDL2 receptors were also less sensitive to GABA compared to *Px*RDL1. Knockout of the fipronil-sensitive *PxRdl1* gene reduced the potency of fipronil by 10 fold, whereas knockout of the fipronil-resistant *PxRdl2* gene enhanced the potency of fipronil by 4.4 fold. Furthermore, in two fipronil-resistant diamondback moth field populations, the expression of *PxRdl2* was elevated by 3.7-fold and 4.1-fold, respectively compared to a susceptible strain, whereas the expression of *PxRdl1* was comparable among the resistant and susceptible strains. Collectively, our results indicate antagonistic effects of *Px*RDL1 and *Px*RDL2 on the fipronil action *in vivo* and suggest enhanced expression of fipronil-resistant *PxRdl*2 potentially a new mechanism of fipronil resistance in insects.

## Introduction

Insect ionotropic GABA receptors are the primary inhibitory neurotransmitter receptor that is widely expressed throughout the insect central nervous system (Sattelle, 1990). The first insect GABA receptor gene was cloned from dieldrin-resistant *Drosophila melanogaster* and designated as *Rdl* (Resistant to dieldrin) (ffrench-Constant *et al*., 1991). RDL GABA receptors belong to the superfamily of pentameric ligand-gated ion channels (LGICs) and thus contain a long N-terminal extracellular domain and four transmembrane regions (TM1-TM4), the second of which (TM2) provides many of the residues that line an integral chloride channel (Casida and Durkin, 2015; Nakao, 2017; Rauh,1990; Ozoe, 2013). Due to their importance in inhibitory transmission, RDL receptors are the target of several classes of highly effective insecticides, such as dieldrin, fipronil and fluralaner (Buckingham *et al*., 2017; Casida and Durkin, 2015; ffrench-Constant *et al*., 2000; Nakao, 2017). Dieldrin belongs to cyclodiene insecticides which are the first generation of noncompetitive antagonists (NCAs) against the RDL receptors (Kadous *et al*.,1983; Rocheleau *et al*.,1993), and fipronil is a second generation NCAs blocking RDL receptors (Gupta and Anadón, 2018; Hosie *et al*., 1995; Zhao *et al*., 2003).

While there is only a single *Rdl* gene in many insect species, such as *D. melanogaster*, *Musca domestica, Apis mellifera* and *Laodelphax striatella* (Eguchi *et al*.,2006; Jones and Sattelle, 2006; Narusuye *et al*., 2007; Rocheleau *et al*., 1993), lepidopteran insects, such as *Plutella xylostella, Bombyx mori, Chilo suppressalis*, and arachnids, such as *Tetranychus urticae* and *Varroa destructor*, possess at least two *Rdl* genes (Dermauw *et al*., 2017; Ménard *et al*., 2018; Sheng *et al*., 2018b; Yu *et al*., 2010; Yuan *et al*., 2010). When expressed in *Xenopus laevis* oocytes, *V. destructor* RDL1 was less sensitive to fipronil than RDL2 and RDL3 (Ménard *et al*., 2018). *C. suppressalis* RDL1 was more sensitive to dieldrin compare to RDL2, but their sensitivity to fipronil was similar (Sheng *et al*., 2018b). To data, three orthologous *Rdl* genes were found from *P. xylostella* (Yuan *et al*., 2010; Zhou *et al*., 2008). However, the importance of individual *Rdl* gene in fipronil action is largely unclear.

It had been reported widely that the alanine to serine or glycine mutation at 2’ position (also known as A2’S or A2’G) and threonine to leucine mutation at 6’ position (also known as T6’L) of RDL GABA receptor are associated with cyclodiene resistance in many insect species such as *D. melanogaster, M. domestica, Haematobia irritans, Blattella germanica* and *Rhipicephalus microplus* (ffrench-Constant *et al*., 1993; Hansen *et al*., 2005; Hope *et al*., 2010; Navarro *et al*., 2010; Thompson *et al*., 1993). Cyclodiene-resistant strains carrying A2’S/G mutations exhibited a low level of cross-resistance to fipronil (Cole *et al*., 1995; Gao *et al*., 2007; Kristensen *et al*., 2005; Nakao, 2017; Scott and Wen 1997). However, a different substitutions at this position, the alanine to asparagine mutation (A2’N) mutation, in RDL has been proved to profoundly decreased the antagonist activity of fipronil in *L. striatellus* and *Sogata furcifera* (Nakao *et al*., 2010; Nakao *et al*., 2011, Sheng *et al*., 2018a). The A2’S mutation in *Px*RDL1 had been reported in *P. xylostella* field strains (Li *et al*., 2006; Wang *et al*., 2016; Yuan *et al*., 2010). When the A2’S mutation was introduced into *Px*RDL1 in an insecticide sensitive *P. xylostella* strain using Clustered Regularly Interspaced Short Palindromic Repeats (CRISPR)-CRISPR-associated protein 9 (Cas9) system, the A2’S mutation caused only about 3-fold increase in fipronil resistance (Guest *et al*., 2019), indicating a limited role of this mutation in fipronil resistance and additional mechanisms underlying higher levels of fipronil resistance in field populations. Interestingly, the *Px*RDL2 has a serine at the 2’position in both susceptible and resistant strains (Jouraku *et al*., 2013; Shi *et al*., 2015; Tang *et al*., 2014; Yuan *et al*., 2010). Whether *Px*RDL2 is involved in fipronil action and resistance, however, remain unknown.

In this study, we evaluated the role of *Px*RDLs in fipronil action and resistance in *P. xylostella* which is one of the most destructive cosmopolitan pests, and has been reported from over 80 countries, and feeds on brassica crops worldwide (Mohan and Gujar, 2003; Sarfraz *et al*., 2005; Zhou *et al*., 2011). Specifically, we compared the transcript of *PxRdl1* and *PxRdl2* between susceptible and resistant populations; and evaluated the sensitivity of *Px*RDL1 and *Px*RDL2 to fipronil in *Xenopus* oocytes. Furthermore, we used the CRISPR-Cas9 technology to knockout *PxRdl1* and *PxRdl2* and evaluated the susceptibility of the resultant mutants to fipronil. Our study revealed distinct roles of *Px*RDL1 and *Px*RDL2 in fipronil action and resistance, and provide valuable information for better understanding the mode of action of fipronil and mechanisms of fipronil resistance.

## Materials and methods

### Insects and chemicals

The susceptible *P. xylostella* strain (He *et al*., 2012) was kindly provided by Dr. Minsheng You (Fujian Agriculture and Forestry University, China). Two field populations were collected from Guangzhou (GZ) city (23.24° N, 113.18° E) of Guangdong province and Fuzhou (FZ) city (26.17° N, 118.51° E) of Fujian province in 2017. The pupae were collected from fields and adults were fed on 10% (V/V) honey solution to lay eggs. The third-instar larvae of the F_1_ strain were subjected to bioassay and total RNA extraction. All populations were maintained separately at 25 ± 1 □, 60-70% relative humidity and under a 16:8 h (light: dark) photoperiod.

In addition to special instructions, all chemicals used in this research were purchased from Sigma-Aldrich (Shanghai, China), and the mature female African clawed frogs (*X. laevis*) were purchased from the Institute of Biochemistry and Cell Biology, SIBS, CAS (Shanghai, China).

### Bioassay

The toxicity of fipronil to *P. xylostella* was tested in leaf-dip bioassays, according to the recommended method of the Insecticide Resistance Action Committee (https://www.irac-online.org). The fipronil was dissolved in dimethyl sulfoxide (DMSO) and diluted with distilled water containing 0.5% Tween-80 to generate five serial dilutions. Leaves of fresh cabbage (*Brassica oleracea*, 2-4 leaf stage) were cut and dipped in insecticide solution for 10 seconds with gentle agitation and placed to surface-dry on paper towel. Control leaves were dipped in distilled water containing 0.5% tween-80 only. 10 third-instar larvae were exposed to the insecticides at each concentration, which was repeated 4 times. Mortality was assessed after 48 h.

### Cloning and sequence of PxRdls

Total RNA was isolated from single third-instar larva of three strains (total 30 larvae of each strain) using the DNA/RNA/Protein Isolation Kit (TianGen, Beijing, China). The first-strand cDNA was synthesized with 1 μg of total RNA using a PrimeScript™ 1st Strand cDNA Synthesis Kit with gDNA Eraser (TaKaRa Biotechnology, Dalian, China). For sequencing analysis of the *PxRdls*, the primers used to amplify the full-length open reading frame (ORF) of *PxRdl1* (GenBank No. NM_001305534.1) and *PxRdl2* (GenBank No. NM_001305535.1) are listed in Table S1. The PCR products of the expected size were purified using the Easy Pure Quick Gel Extraction Kit (Transgen Biotech, Beijing, China) and cloned into TA Vector (Takara Biotechnology, Dalian, China). Positive clones were selected and sequenced by Beijing Genomics Institute (Beijing, China). DNA alignments were performed by DNAman V9 (Lynnon Biosoft, CA, USA).

### Expression analysis of PxRdls mRNA

To monitor the transcript level of *PxRdls*, Total RNA was extracted from 5-6 third-instar larvae and repeated 3 times. qRT-PCR was carried out using the SYBR^®^ Premix Ex Taq™ II (Takara Biotechnology, Dalian, China) and CFX96 Connect Real-Time system (Bio-RAD, USA). The qRT-PCR program was 95 °C for 2 min, followed by 40 cycles of 95 °C for 10 s, 60 °C for 30 s and 72 °C for 30 s. Melting curves were prepared by increasing the temperature from 60 to 95 °C to check the specificity of the primers when the cycling reaction finished. qRT-PCR data was normalized to an internal control, the *Actin* gene (GenBank No. JN410820.1), and analyzed using the 2^−ΔΔCT^ method.

### cRNA preparation and X. laevis oocytes injection

The ORFs encoding *PxRdls* were cloned into a pT7TS vector using the In-Fusion HD Cloning Kits (Takara Biotechnology, Dalian, China) and, finally validated by sequencing (Fig. 1). The linearized recombinant plasmid was used as templates to synthesize capped RNAs (cRNAs) with the mMessage mMachine T7 Kit (Life Technologies, Carlsbad, CA). The cRNAs were dissolved in RNase-free water at a concentration of 1000 ng/μL, and finally stored at −80□.

**Fig.1.**
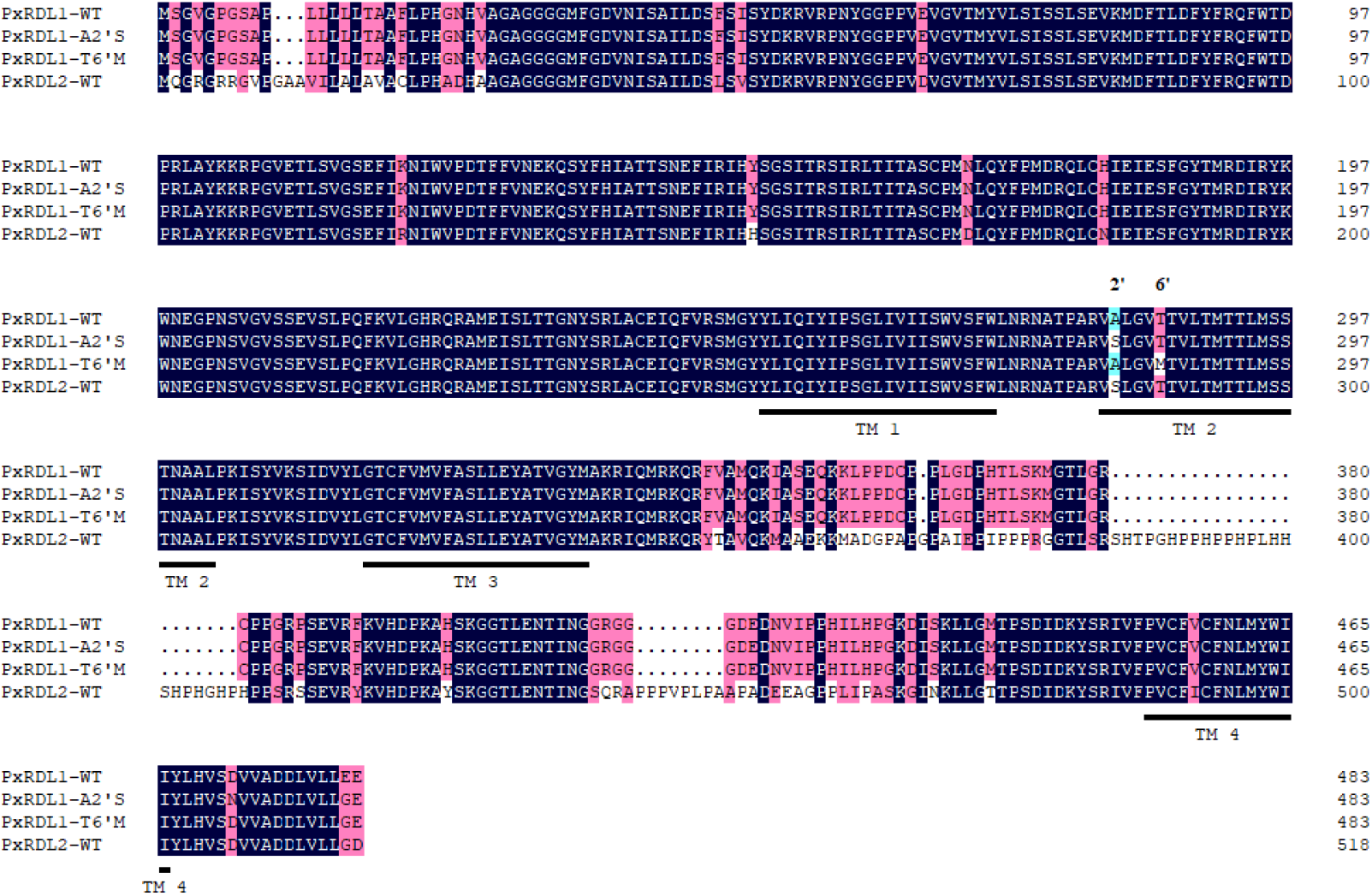
The alignment of amino acid sequence differences between *Px*RDL1-WT, *Px*RDL1-A2’S, *Px*RDL1-T6’M and *Px*RDL2-WT. WT, the wild type *Px*RDL1 and *Px*RDL2; A2’S, the alanine to serine mutation in *Px*RDL1; T6’M, the threonine to methionine mutation in *Px*RDL1. The shown sequences also were the sequences that expressed in *Xenopus laevis* oocytes.

Mature female frogs were anesthetized in 1 g/L ethyl 3-aminobenzoate methanesulfonate for 15-20 min. *X. laevis* oocytes at stage □ and □ were harvested and rinsed with Ca^2+^-free SOS solution (100 mM NaCl, 2 mM KCl, 1 mM MgCl_2_, 5 mM 4-(2-hydroxyethyl)-1-piperazineethanesulfonic acid (HEPES), pH 7.6), and then treated with 1 mg/ mL type □ collagenase in Ca^2+^ free SOS solution. After 1-2 h of incubation at 16□, the oocytes were gently washed with SOS solution (100 mM NaCl, 2 mM KCl, 1 mM MgCl_2_, 1.8 mM CaCl_2_, 5 mM HEPES, pH 7.6) containing 2.5 mM sodium pyruvate, 50 mg/ml gentamycin 50 mg/ ml, 100 U/ml penicillin and 100 mg/ml streptomycin, and then incubated in the same solution at 16□ overnight. The oocytes were injected with 4.6 ng of cRNA. The injected oocytes were incubated in SOS solution at 16□ for electrophysiological experiments.

### Two-electrode voltage-clamp (TEVC) recordings

Oocytes 2-4 days post-injection were kept in SOS solution and clamped at −80 mV. The currents were recorded by the two electrode voltage-clamp (TEVC) method using an Oocyte Clamp OC-725C amplifier (Warner Instruments, Hamden, CT, USA) with continuous perfusion at a rate of 3 mL/min. The recording pipettes were filled with 3 M KCl, delivering an electrical resistance of 0.5-2 MΩ. GABA was dissolved in SOS solution, and fipronil was dissolved in DMSO and then diluted with SOS solution with a final DMSO concentration of less than 0.1% (v/v). GABA concentration response curves were obtained by perfusing oocytes with GABA concentrations from 1 μM to 1 mM. Responses to GABA at each concentration were normalized to the maximum current induced by 1 mM GABA. Fipronil inhibition curves were generated by inhibiting the response to GABA (EC50). The oocytes were first stimulated with GABA (EC_50_) twice, followed by perfusing the fipronil alone for 5 min, and finally GABA (EC_50_) was co-applied with fipronil. Each experiment was performed on 5-6 different oocytes obtained from at least two frogs.

### sgRNA preparation and eggs microinjection

The single guide RNA (sgRNA) for editing *PxRdl1* and *PxRdl2* gene were designed using CRISPR RGEN Tools (http://www.rgenome.net/). After the off-target analysis, the sgRN*A-PxRdl1* target sequence (5’ TTAGCGTATAAAAAAAGGCC*AGG* 3’) was selected at exon 4 of *PxRdl1* (Fig. 5A), and the sgRN*A-PxRdl2* target sequence (5’*CCT*GGCGCTGCCGTCATCCTGGC 3’) was selected at exon 1 of *PxRdl2* (Fig. 6A). The sgRNAs were prepared following the instruction (Guide-it™ sgRNA *In Vitro* Transcription and Screening Systems, TaKaRa, Japan). Briefly, SgRNA synthesis templates were amplified firstly, and the PCR product was run and analyzed on 2% agarose gel to confirm the size of the PCR product is about 130bp. Then combine 20 μL *in vitro* transcription reaction solution contain 5 μL of PCR product, 7μL of Guide-it *In Vitro* Transcription Buffer, 3 μL of Guide-it T7 Polymerase Mix and 5μL of RNase Free Water. The transcription reaction solution was incubated at 37°C for 4 h to maximize sgRNA yield. Following incubation, 2 μL of DNase I was added and incubated at 37□ for 15 min. Finally the sgRNA was purified with Guide-it IVT RNA Clean-Up Kit (TaKaRa, Japan) and stored at −80□ for injection step.

Microinjection of Cas9 protein and sgRNA was conducted as the method described by Huang *et al*., (2016). In brief, susceptible strains of female *P. xylostella* were allowed to lay eggs on parafilm sheets treated previously with cabbage juice. The fresh eggs were collected within 30 min and microinjected immediately with the mixture of 300 ng/μL Cas9-N-NLS Nuclease (GenCrispr, China) and 150 ng/μL sgRNA. The Cas9 protein and sgRNA were mixed well and incubated at 37□ for 15 min before injection. Injection was carried out using an Eppendorf TransferMan^®^ 4r micromanipulator, Eppendorf FemtoJet^®^ 4x programmable microinjector and Eppendorf Femtotip II injection capillaries (Eppendorf, Hamburg, Germany). The injection conditions were injection pressure (p_i_) 600 hPa, compensation pressure (p_c_) 80 hPa and injection time (t_i_) 0.8s. Eggs after injection were incubated at 25 ± 1□, 60-70% relative humidity for hatching.

### Identification of PxRdl mutants

About 500 fresh eggs were injected with the mixture of Cas9 and sgRNA. Finally, 43 and 37 of the eggs injected with sgRNA-*PxRdl1* or sgRNA-*PxRdl2* successfully developed to moths (named as G_0_ progeny). Single-pair mating between G_0_ and wild type adults were set up to generate G_1_ progeny.

The G_0_ progeny were collected after oviposition, and the genomic DNA was extracted individually using TIANamp Genomic DNA Kit (TianGen, Beijing, China). The detection primers (Table S1) were designed to amplify the target region flanking the desired cleavage site. The PCR product was directly sequenced to identify the detailed indels of genomic DNA. qRT-PCR was performed to measure the transcript level of *PxRdls* in homozygous mutant strains.

### Statistical analysis

The bioassay, qRT-PCR and electrophysiology data are presented as the means ± standard error (SE). For the bioassay, the LC_50_ of fipronil was obtained by a linear regression program using SPSS 22.0 (SPSS Inc., IL). For electrophysiology analyses, the EC_50_ and IC_50_ were obtained by a nonlinear regression program using GraphPad Prism 5.0 (GraphPad Software Inc., La Jolla, CA, USA). The mRNA expression significance was analyzed with one-way ANOVA followed by Tukey’s test (*p* < 0.01) using SPSS 22.0 (SPSS Inc., IL). The significance of LC_50_, EC_50_ and IC_50_ were analyzed with Student’s *t*-test (alpha=0.05) using Stata v. 14 (StataCorp LLC, College Station, TX, USA).

## Results

### Resistance to Fipronil

Fipronil toxicity analysis was performed on the two field (GZ and FZ) F1 strains and the susceptible strain by leaf-dip bioassays. The two field strains exhibited high levels of fipronil resistance, with resistance ratios that were 1686-fold for the GZ populations and 953-folds for FZ populations relative to the susceptible strain (Table 1).

**Table 1.** Resistance to fipronil in field-collected strains of *P. xylostella*.

| Strain | N<sup>a</sup> | LC<sub>50</sub> (95% CI) (mg L<sup>-1</sup>)<sup>c</sup> | Slope ± SE | RR<sup>b</sup> |
| --- | --- | --- | --- | --- |
| Susceptible | 237 | 0.29 (0.24-0.34) a | 1.94±0.19 |  |
| GZ | 241 | 488.97 (402.81-605.28) c | 1.65±0.18 | 1686 |
| FZ | 236 | 276.32 (222.30-332.66) b | 1.80±0.19 | 953 |

<sup>a</sup>Numbers of larvae used in bioassay.
<sup>b</sup>RR(resistance ratio)=LC<sub>50</sub> ( GZ or FZ ) /LC<sub>50</sub>(Susceptible) .
<sup>c</sup> Lower case letters(a, b and c) indicate significance between different strains. Statistical analysis was determined by Student's *t*-test (alpha=0.05)

### Mutation screening of PxRDLs in field resistant populations

Results from sequence analysis showed that, for *Px*RDL1, compared with the amino acid sequence from the susceptible strain, A2’S mutation was present in the GZ population with a frequency of 73.33%, while the frequency was 33.33% in the FZ population. Furthermore, another mutation, T6’M, was detected in the GZ population with a lower frequency of 23.33% and higher frequency of 56.67% in the FZ population (Fig.1 and Table 2). Individuals carrying A2’S and T6’M double mutations in *Px*RDL1 were not found in any of the populations examined. There were only few mutations with low frequency (3.33%-18.33%) been detected in the N-terminal extracellular domain and TM3-TM4 loop of *Px*RDL2 (Table S2).

**Table 2.**
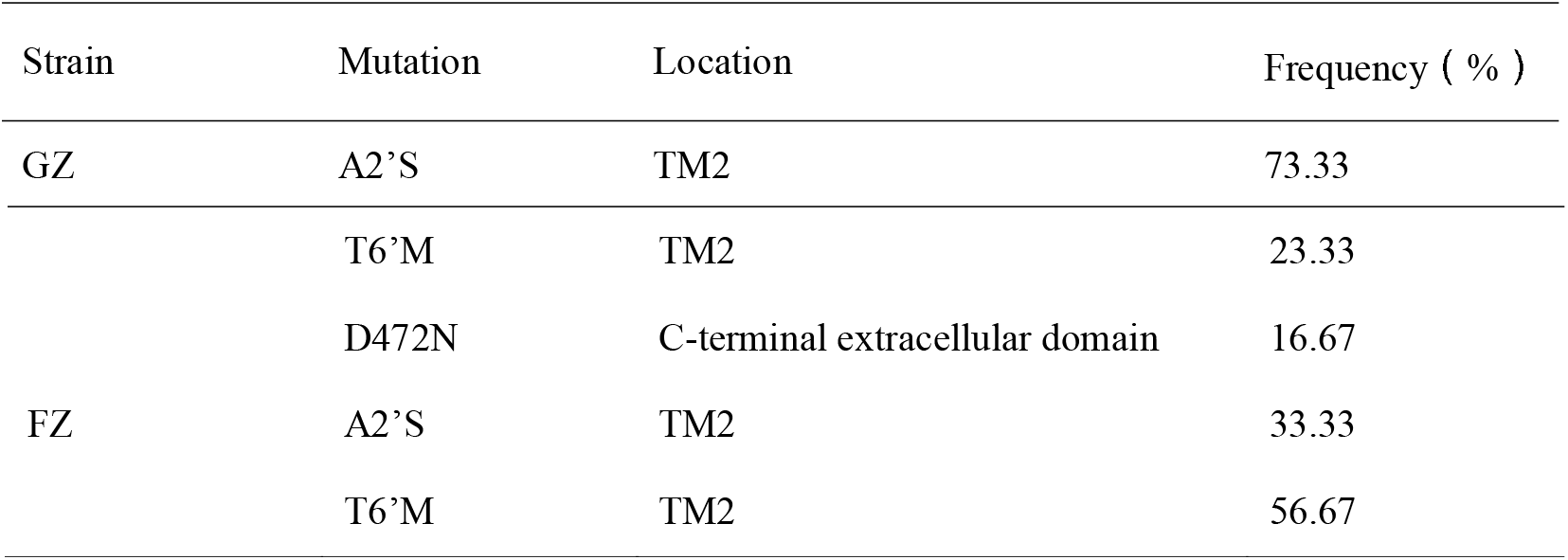
The amino acid mutation frequencies of the *Px*RDL1 subunit in the GZ and FZ strains.

### Functional and pharmacological characterization of PxRDLs in Xenopus oocytes

To compare the sensitivity of the *Px*RDLs to fipronil, *Px*RDLs were expressed in *Xenopus* oocytes, and the potencies of fipronil as an antagonist and GABA as an agonist were evaluated under the TEVC condition. The EC_50_ value of GABA in activating currents in oocytes expressing *Px*RDL2 was 153.40 μM, which was ≈ 7-fold higher than that in oocytes expressing *Px*RDL1 (Fig. 3; Table 3). The IC_50_ for fipronil in oocytes expressing *Px*RDL2 was much higher than that in oocytes expressing *Px*RDL1, with an IC_50_ value of 5.23 μM (Fig. 4; Table 3).

**Table 3.**
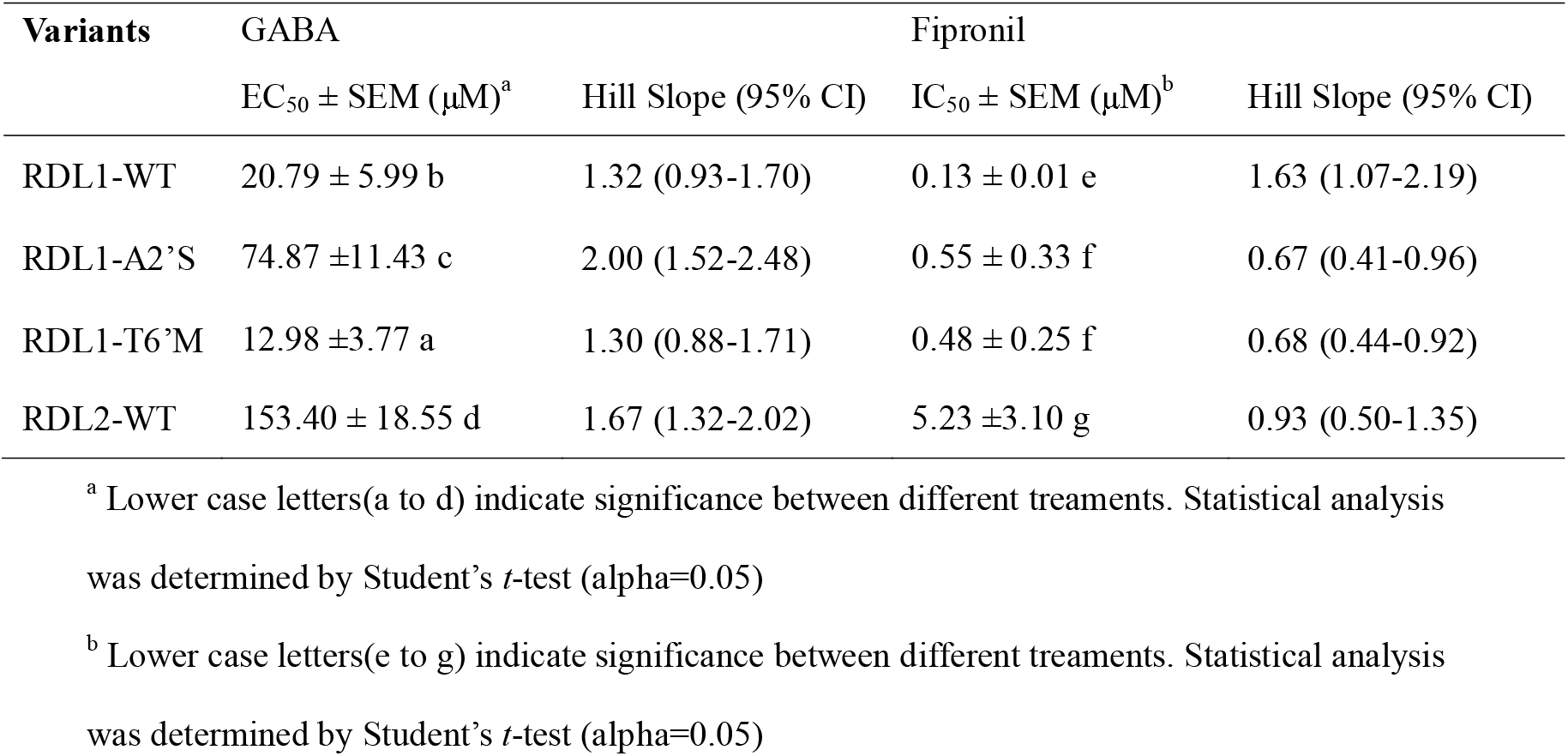
Potencies of GABA and fipronil in *Xenopus* oocytes injected with *PxRdl* cRNAs

### Over-expression of PxRDL2 transcript in the resistant strains

qRT-PCR was performed to compare the expression of the *PxRdls* genes among the susceptible and two resistant strains. The results showed that the transcript level of *PxRdl1* was 6.92-fold higher than that of *PxRdl2* in the susceptible strain (Fig. 2A). There was no significant difference in the transcript level of the *PxRdl1* among the three strains. However, the transcript level of *PxRdl2* was 4.11-fold higher in the GZ strain and 3.70-fold higher in the FZ strain compared to the susceptible strain (Fig. 2A).

**Fig.2.**
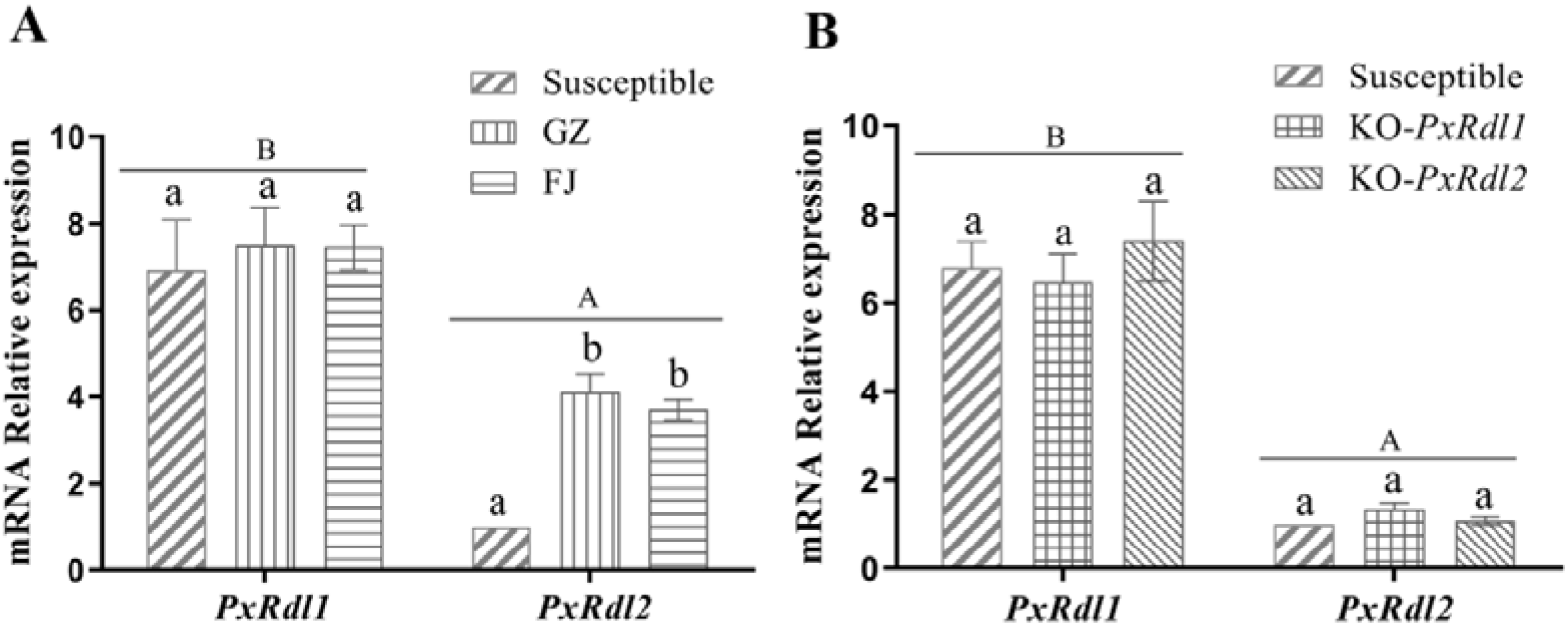
The expression profile of *PxRdls* mRNA in third-instar larvae of different strains. (A) The mRNA expression level of susceptible and two filed strains. (B) The mRNA expression level of susceptible and two *PxRdls* knockout strains. Error bars indicate the SE of the mean of three independent replicates. Uppercase letters (A and B) indicate significance between *PxRdl1* and *PxRdl2* in the same strain. Lower case letters (a and b) indicate significance among different strains. The significance was analyzed with one-way ANOVA followed by a Tukey test (*P* <0.01).

### Construction of homozygous PxRdl mutant P. xylostella strains

To further evaluate the role of *Px*RDL2 in the action of fipronil, we used CRISPR-Cas9 to construct *KO-PxRdl1* and *KO-PxRdl2* strains, respectively. G_1_ progeny of *PxRdl1* mutants were obtained and multiple deletion types were found in target sequence, such as 4-nt deletion (Fig. 5B) and 2, 3, 12, 13-nt deletion (data not shown). There were also multiple deletion types, such as 1, 6, 18-nt deletion (data not shown) and a 52-nt deletion near the target sequence been detected in the G_1_ progeny of *PxRdl2* mutants (Fig. 6B).

The 4-nt deletion of *PxRdl1* (7 pairs) and 52-nt of *PxRdl2* (9 pairs) heterozygous mutant G_1_ adults were set up single-pair mating to obtain G_2_ progeny. 62 pairs of 4-nt deletion of *PxRdl1* and 70 pairs of 52-nt deletion of *PxRdl2* adults were set up single-pair mating to obtain G_3_ progeny, respectively. Then the single-pair mating was performed progeny by progeny, until the male and female adults were both homozygous mutant (Fig. S1). Genomic DNA sequencing of G_2_ progeny adults confirmed that 5 pairs of 4-nt deletion of *PxRdl1* and 4 pairs of 52-nt deletion of *PxRdl2* adults were homozygous mutant. All adults of the G_3_ progeny generated from homozygous mutant G_2_ progeny were still set up single-pair mating and genomic DNA sequencing to confirm they were homozygous mutant. In conclusion, we successfully obtained a homozygous mutant strain (KO-*PxRdl1*) that has 4-nt deletion at exon 4 of *PxRdl1*, and a homozygous mutant strain (KO-*PxRdl2*) that has 52-nt deletion at exon 1 of *PxRdl2*. The 4-nt and 52-nt deletion both resulted in frame shift and premature stop codons (Fig. 5D and Fig. 6D).

### Impact of PxRDL disruption on fipronil toxicity

qRT-PCR results showed that there was no significant difference in individual *PxRdl* transcript level between wild-type and *PxRdl* knockout strains (Fig. 2B). Bioassay results showed that compared to the wild-type strain, *KO-PxRdl1* larvae exhibited 10.41-fold fipronil resistance; however, *KO-PxRdl2* larvae were 4.4-fold more sensitive to fipronil (Table 4).

**Table 4.**
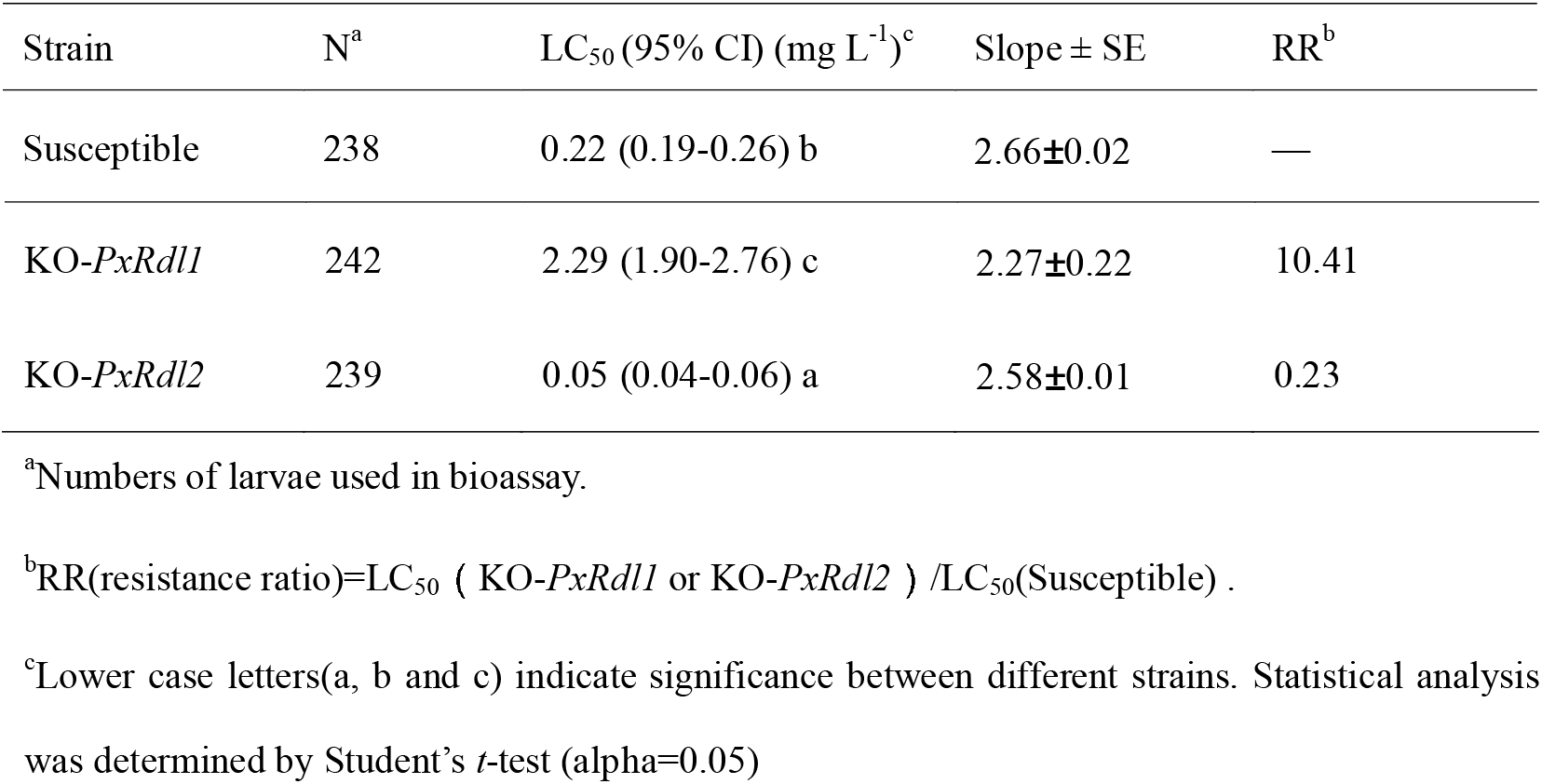
Toxicity of fipronil to susceptible and *PxRdl* knockout *P. xylostella*.

| Strain | N<sup>a</sup> | LC<sub>50</sub> (95% CI) (mg L<sup>-1</sup>)<sup>c</sup> | Slope ± SE | RR<sup>b</sup> |
| --- | --- | --- | --- | --- |
| Susceptible | 238 | 0.22 (0.19-0.26) b | 2.66±0.02 | — |
| *KO-PxRdl1* | 242 | 2.29 (1.90-2.76) c | 2.27±0.22 | 10.41 |
| *KO-PxRdl2* | 239 | 0.05 (0.04-0.06) a | 2.58±0.01 | 0.23 |

<sup>a</sup>Numbers of larvae used in bioassay.
<sup>b</sup>RR(resistance ratio)= $LC_{50}$ ( *KO-PxRdl1* or *KO-PxRdl2* ) / $LC_{50}$ (Susceptible) .
<sup>c</sup>Lower case letters(a, b and c) indicate significance between different strains. Statistical analysis was determined by Student's *t*-test (alpha=0.05)

## Discussion

In this study, we evaluated the roles of *Px*RDL1 and *Px*RDL2 in fipronil action and resistance. Our functional expression of the *Px*RDL receptors in *Xenopus* oocytes showed that the *Px*RDL1 monomeric receptor is significantly more sensitive to fipronil than the *Px*RDL2 monomeric receptor. Our CRISPR-Cas9 experiments showed that *Px*RDL1 knockout insects are more resistant to fipronil, whereas *Px*RDL2 knockout insects are more sensitive to fipronil. Furthermore, besides detection of two previously identified A2’S and T6’M mutations in *Px*RDL1 from two fipronil-resistant field strains, we discovered an elevated level of *Px*RDL2 transcripts (but not *Px*RDL1 transcripts) in both strains. Our results suggest an antagonistic effect of *Px*RDL2 on the action of fipronil and elevated expression of *Px*RDL2 as a potential new mechanism of fipronil resistance.

Earlier studies reported that *P. xylostella* possess three orthologous *Rdl* genes. *Px*RDL1 and *Px*RDL3 (GenBank No. EF156251) both have an amino acid alanine at 2’ position, and *Px*RDL2 has a 2’ serine (Yuan *et al*., 2010; Zhou *et al*., 2008). We failed to amplify *Px*RDL3 by RT-PCR although we attempted using various PCR primer pairs. Therefore, in this study we evaluated the sensitivity of *Px*RDL1 and *Px*RDL2 to fipronil in *X. laevis* oocytes. *Px*RDL1 and *Px*RDL2 exhibited different sensitivities to both fipronil and GABA (Figs. 3 and 4). *Px*RDL2 was 40.23-fold less sensitive to firponil that *Px*RDL1 (Fig. 4). *Px*RDL2 is also less sensitive to GABA compared to *Px*RDL1 (Fig. 3). These results are consistent with those from a functional analysis of RDLs from *V. destructor* (Ménard *et al*., 2018). Like *Px*RDL1, *V. destructor* RDL2 and RDL3 monomeric receptors have an amino acid alanine at 2’ position and are more sensitive to fipronil and GABA. Like *Px*RDL2, the *V. destructor* RDL1 monomeric receptor has 2’ serine and exhibits a lower fipronil sensitivity and GABA affinity (Ménard *et al*., 2018). However, *C. suppressalis* RDL1 and RDL2 which have 2’ alanine and 2’ serine, respectively, display a similar sensitivity to fipronil (Sheng *et al*., 2018b). Nevertheless, *C. suppressalis* RDL1 was more sensitive to dieldrin than RDL2 (Sheng *et al*., 2018b). These results suggest that additional unidentified sequence features contribute to the low sensitivity to fipronil of *V. destructor* RDL1 and *Px*RDL2 receptors.

**Fig.3.**
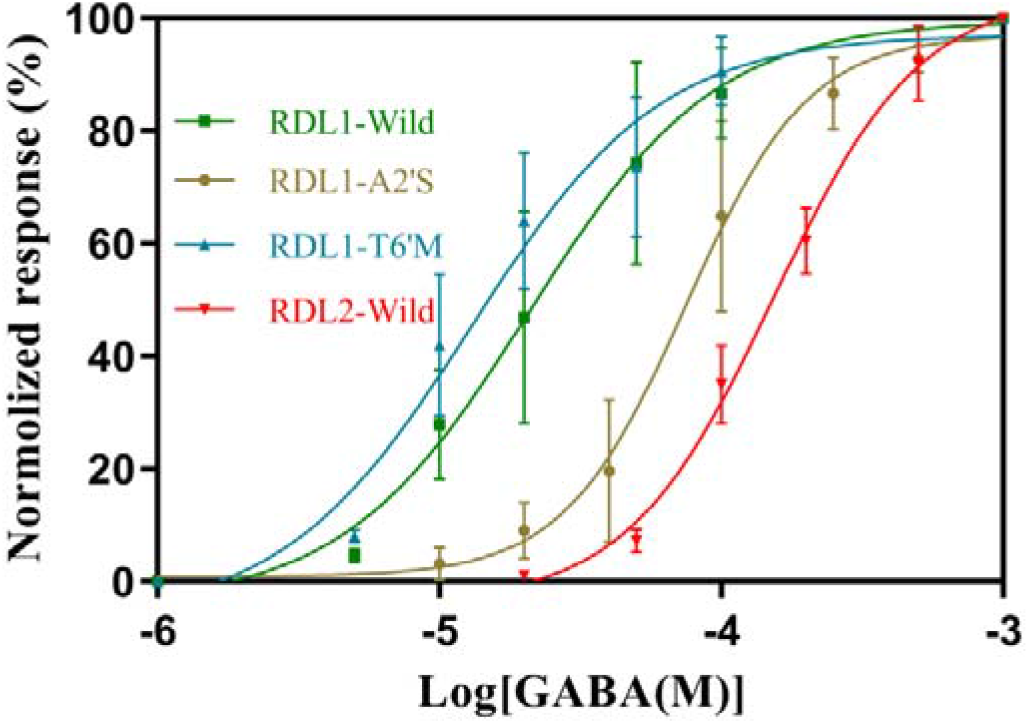
Dose-response curves of GABA-induced current in oocytes injected with *PxRdl* cRNAs. Each point represents the mean ± SE of responses in 5-6 oocytes from at least two *Xenopus* laevis.

**Fig.4.**
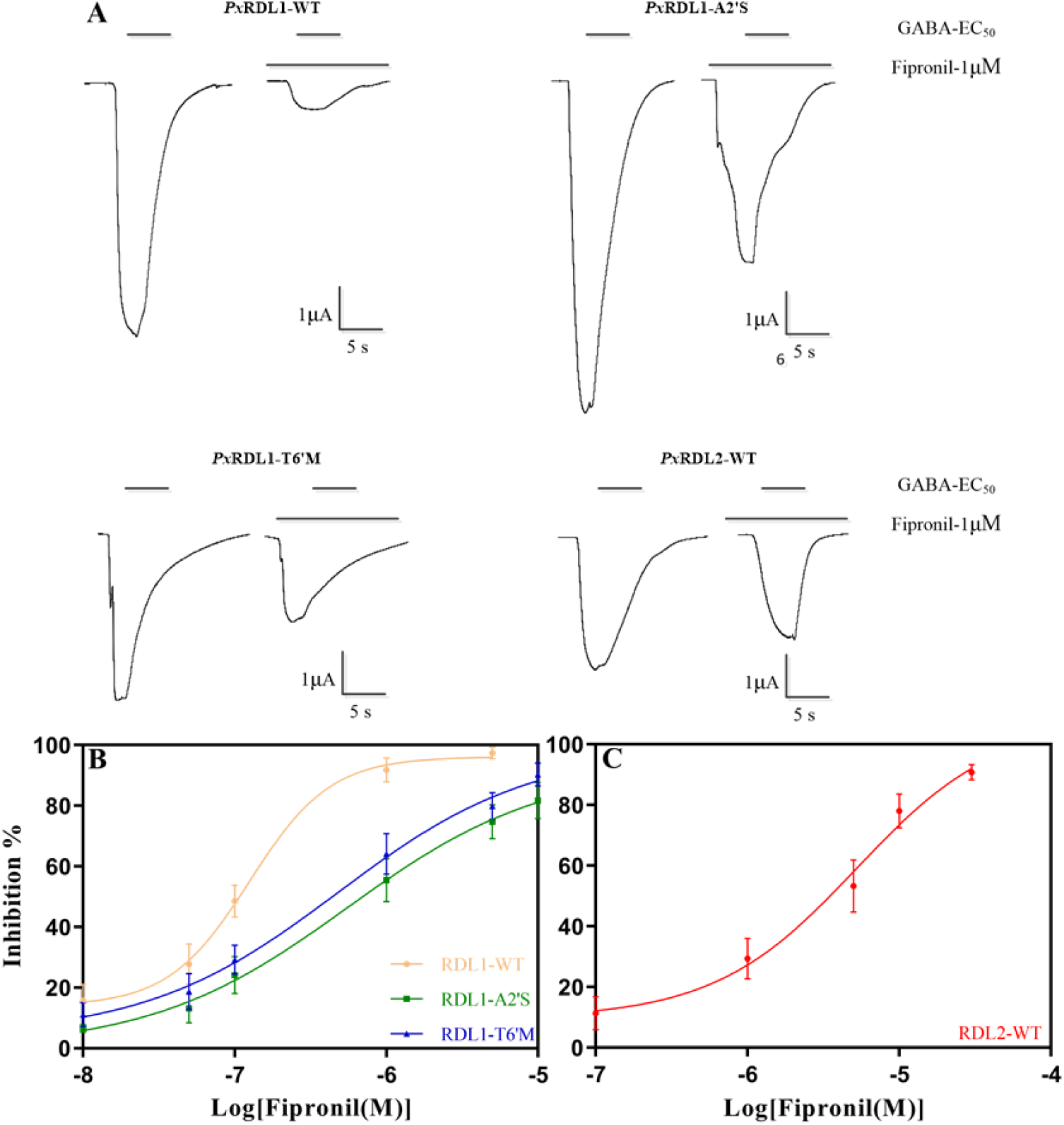
Fipronil inhibition of GABA-induced currents in oocytes injected with *PxRdl* cRNAs. (A) Examples of fipronil inhibition of currents in channels containing *PxRdl* cRNAs (B) Fipronil concentration-response curves in channels containing *Px*RDL1. (C) Fipronil concentration-response curves in channels containing *Px*RDL2. Each point represents the mean ± SE of responses in 5-6 oocytes from at least two *Xenopus* laevis.

**Fig.5.**
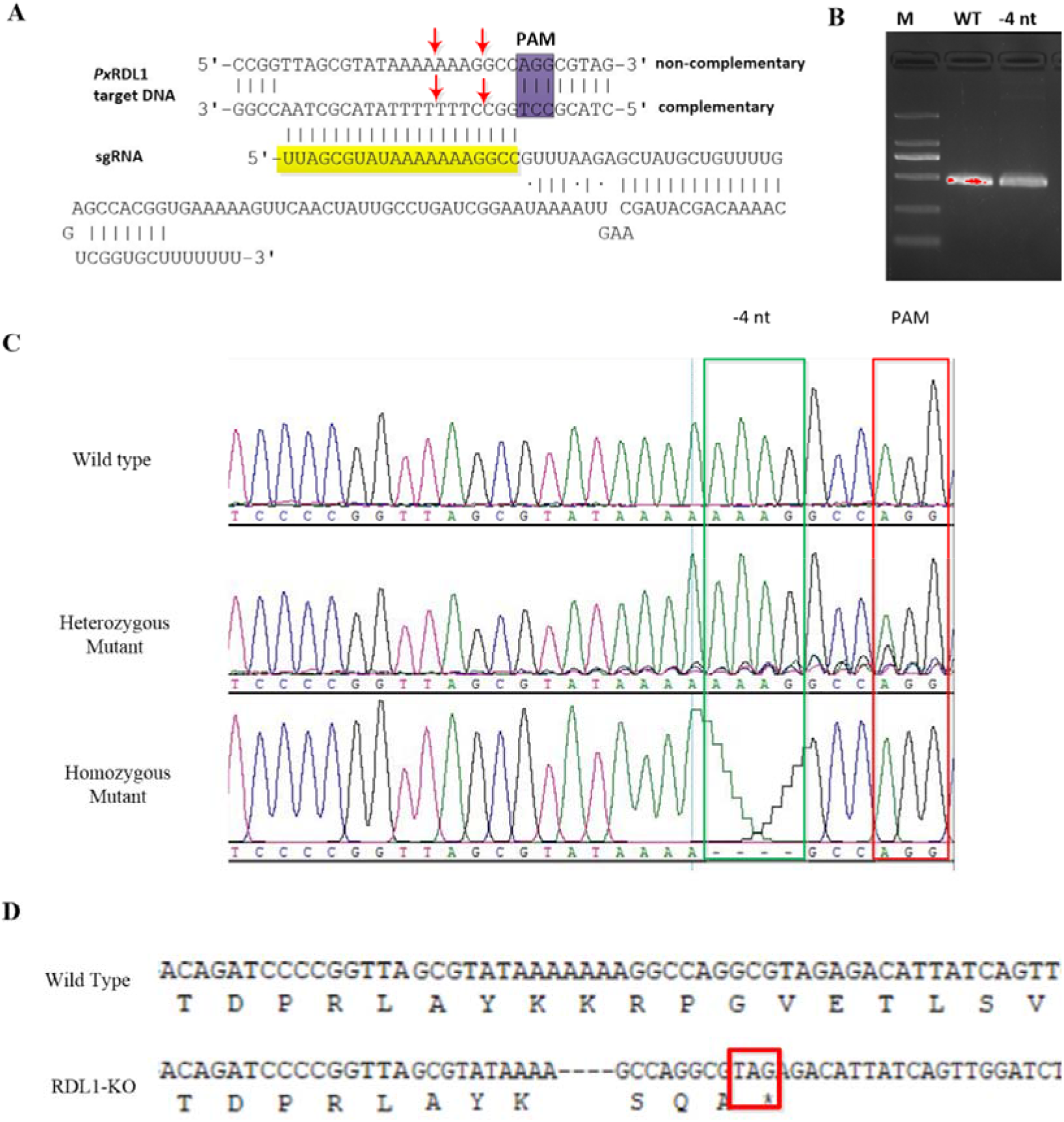
The CRISPR-Cas9 mediated mutation type of *PxRdl1* gene. (A) Schematic representation of sgRNA and *PxRdl1* DNA sequences. The protospacer DNA is shown in yellow and protospacer adjacent motif (PAM) sequence is shown in purple. Cleavage sites are represented by red arrows. (B) PCR was performed using primers designed to detect the 4-nt deletiion in *PxRdl1*. M, marker. WT, wild type genomic DNA. (C) Direct sequencing chromatograms of PCR products amplified from genomic DNA of wild type, heterozygous mutant and homozygous mutant (−4nt deletion). The locations of −4nt deletion and PAM were marked by green and red box, respectively. (D) The deduced peptide sequence from partial of exon 4 of *PxRdl1*, the stop code was marked by red box.

**Fig.6.**
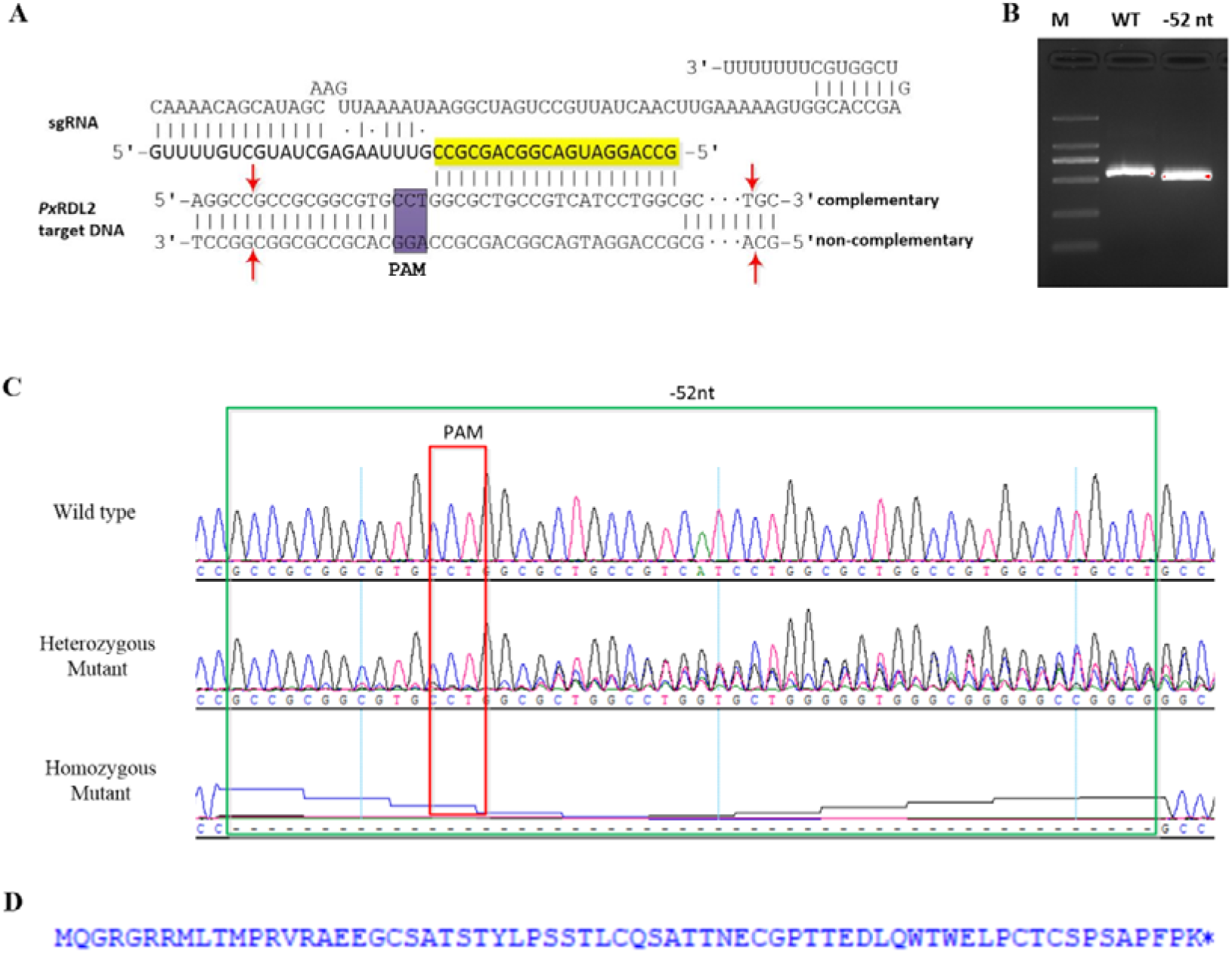
The CRISPR-Cas9 mediated mutation type of *PxRdl2* gene. (A) Schematic representation of sgRNA and *PxRdl2* DNA sequences. The protospacer DNA is shown in yellow and protospacer adjacent motif (PAM) sequence is shown in purple. Cleavage sites are represented by red arrows. (B) PCR was performed using primers designed to detect the 52-nt deletion in *PxRdl2*. M, marker. WT, wild type genomic DNA. (C) Direct sequencing chromatograms of PCR products amplified from genomic DNA of wild type, heterozygous mutant and homozygous mutant (−52nt deletion). The locations of −52nt deletion and PAM were marked by green and red box, respectively. (D)The deduced peptide sequence of *Px*RDL2, the 52nt deletion caused premature translation termination of *Px*RDl.2.

Wei *et al*., (2015) knocked down the transcript level of the *L. striatellus Rdl* gene using RNA interference (RNAi), and found the effect of fipronil was significantly reduced. Similarly, Jia *et al*., (2019) also found that knockdown the transcript level of *Rdl1* or *Rdl2* gene both significantly reduce the mortality of *C. suppressalis* treated with fluralaner (mainly acting on RDL GABAR). CRISPR-Cas9 technology has been successfully applied in insects to clarify the relationship between targets and insecticide action *in vivo* (Homem and Davies, 2018; Wang *et al*., 2017), as it can block the gene function completely. For instance, Wang *et al*., (2020) proved the *P. xylostella* nicotinic acetylcholine receptors (nAChRs) α6 gene is the main target of spinosyns, and knockout of this gene caused over 200 fold spinosyns resistance. In this study, our results revealed distinct roles of *Px*RDLs in the action and resistance of fipronil in *P. xylostella*. Knockout of *PxRdl1* gene conferred resistance to fipronil, and *Px*RDL1 receptors are highly sensitive to fipronil, confirming that *Px*RDL1 is a main target of fipronil. However, knockout of fipronil-resistant *PxRdl2* gene enhanced insect sensitivity to fipronil, revealing an antagonistic effect of *Px*RDL2 on the action of fipronil in *P. xylostella*. Our qRT-PCR results showed that the transcript level of *PxRdl1* did not change in the *PxRdl2* knockout strain, and *vice versa* (Fig. 2B), suggesting no obvious genetic compensation at the transcriptional level. Furthermore, knockout of *PxRdl1* and *PxRdl2* didn’t induce lethality. These results provide evidence for functional redundancy regarding the role of *Px*RDL1 and *Px*RDL2 in neuronal signaling, despite their distinct contributions to the action of fipronil.

The presence of multiple *Rdl* genes was thought to be the result of gene duplication or conversion events that occurred recently to enhance the tolerance to naturally occurring toxins or insecticides (Ménard *et al*., 2018; Meng *et al*., 2019; Sheng *et al*., 2018b). In this study, we showed that *Px*RDL2 was more resistant to fipronil than *Px*RDL1. We speculate that the increased transcript level of *PxRdl2* gene in the two fipronil resistant field strains may be associated with fipronil resistance. Furthermore, although how *Px*RDL2 antagonizes the action of fipronil remains to be determined, enhanced level of *PxRdl2* transcripts could impose a greater antagonism on the action of fipronil, thereby, confer resistance to fipronil.

## Acknowledgment

We would like to thank Dr. Youming Sheng and Dr. Weiyi He for their helping in knockout genes in *P. xylostella* by CRISPR-Cas9 technology. The work was supported by the National key R&D program of China (2018YFD0200300), the Project of Science and Technology in Guangdong Province (2018A030313188), the Natural Science Foundation of Guangdong Province (2017A030310490) and the Research and Innovation Team of Key Technologies in Modern Agricultural Industry in Guangdong Province (2019KJ130).

