## Supplementary material for "Distinct roles of two RDL GABA-receptors in fipronil action in the diamondback moth (*Plutella xylostella*)": supplemetal Table1 and 2, supplemental Fig 1

**Table S1** Detailed information of primers used in this study.

| Primer name | Nucleotide sequence (5´-3´) | Function |
| --- | --- | --- |
| *Rdl1*-F | GCGCCAGCCCGCTCCCATG | Amplification of *PxRdl1* |
| *Rdl1*-R | CTATTTATCCTCCCCCAGAAGCACC |  |
| *Rdl2*-F | TTGTAGTGCGTCCGTTAGGGCG | Amplification of *PxRdl2* |
| *Rdl2*-R | GATGGTCATTGCCCTAGTTTCA |  |
| qPCR-*Rdl1-*F | GGCAATCATGTCGCGGGTGCAGGTG | Quantitative real-time PCR |
| qPCR-*Rdl1*-R | ATCCATTTTCACTTCAGACAGAGAG |  |
| qPCR- *Rdl2*-F | GTCAGTCAGCTACGACAAACG |  |
| qPCR- *Rdl2*-R | AAATCCAGGGTAAAATCCAT |  |
| qPCR-Actin-F | TGGCACCACACCTTCTAC | Reference gene |
| qPCR-Actin-R | CATGATCTGGGTCATCTTCT |  |
| pT7-*Rdl1*-F^a^ | **AGATCTGATATCACTAGT***GCCACC*ATGAGCGGCGTCGGCCCGGGCAGC | Amplification of ORF for oocytes expression |
| pT7-*Rdl1*-R^a^ | **CGCGGCCGCCTCGAGGCATGC**CTATTTATCCTCCCCCAGAAGCACC |  |
| pT7-*Rdl2*-F^a^ | **AGATCTGATATCACTAGT***GCCACC*ATGCAAGGCAGAGGCCGCCGC |  |
| pT7-*Rdl2*-R^a^ | **CGCGGCCGCCTCGAGGCATGC**TTAGTTAATGTCTCCAAGTAGTACC |  |
| Crispr-*Rdl1*-F^b^ | CCTCTAATACGACTCACTATAGG*TTAGCGTATAAAAAAAGGCC*GTT  TAAGAGCTATGC | sgRNA synthesis template sequence amplification |
| Crispr-*Rdl2*-F^b^ | CCTCTAATACGACTCACTATAG*GCCAGGATGACGGCAGCGCC*GTTT  AAGAGCTATGC |  |
| Detection-*Rdl1*-F | TCATTGCTCCAAACTG | gDNA detection |
| Detection-*Rdl1*-R | CTTCTGGTAATAGACCCT |  |
| Detection-*Rdl2*-F | CTTCGGTGGGAGATGG |  |
| Detection-*Rdl2*-R | TACAGTTGTGCGACGAGA |  |

Note: ^a^The bold nucleotide sequences are infusion sequence used to connect to the pT7TS vector. The italic nucleotide sequence *GCCACC* is the Kozak sequence.

^b^ The N_20_ sequence was Italicized.

**Table S2** The amino acid mutation frequencies of *Px*RDL2 in GZ and FZ strain.

| Strain | Mutation | Location | Frequency（%） |
| --- | --- | --- | --- |
| GZ | P10S | N-terminal extracellular domain | 6.67 |
|  | A13V | N-terminal extracellular domain | 10.00 |
|  | A32T | N-terminal extracellular domain | 13.33 |
|  | V111I | N-terminal extracellular domain | 3.33 |
|  | R220M | N-terminal extracellular domain | 10.00 |
|  | G223S | N-terminal extracellular domain | 6.67 |
|  | S239L | N-terminal extracellular domain | 3.33 |
|  | S401N | TM3–TM4 loop | 10.00 |
|  | T435I | TM3–TM4 loop | 13.33 |
| FZ | A32T | Membrane-spanning region 2 | 6.67 |
|  | D68E | Membrane-spanning region 2 | 16.67 |
|  | L81V | N-terminal extracellular domain | 13.33 |
|  | K85L | N-terminal extracellular domain | 18.33 |
|  | S118P | N-terminal extracellular domain | 16.67 |
|  | S212C | N-terminal extracellular domain | 13.33 |
|  | S250P | N-terminal extracellular domain | 6.67 |
|  | P373H | TM 3–TM 4 loop | 10.00 |
|  | H404R | TM 3–TM 4 loop | 3.33 |
|  | S412T | TM 3–TM 4 loop | 6.67 |
|  | S427P | TM 3–TM 4 loop | 10.00 |
|  | P447S | TM 3–TM 4 loop | 6.67 |
|  | I468T | TM 3–TM 4 loop | 3.33 |
|  | K470E | TM 3–TM 4 loop | 8.33 |

**Fig.S1** Flow diagram of the crossing scheme used to obtain the *PxRdl1* and *PxRdl2* homozygous knockout strains. The injected eggs were named as G_0_ progeny. Single-pair mating between G_0_ and wild type adults were set up to obtain G_1_ progeny. G_1_ adults were set up single-pair mating to obtain G_2_ progeny, followed by genomic DNA sequencing. G_2_ progeny generated by mutant G_1_ adults were collected and set up single-pair mating. Then the single-pair mating was performed progeny by progeny, until the male and female adults were both homozygous mutant.

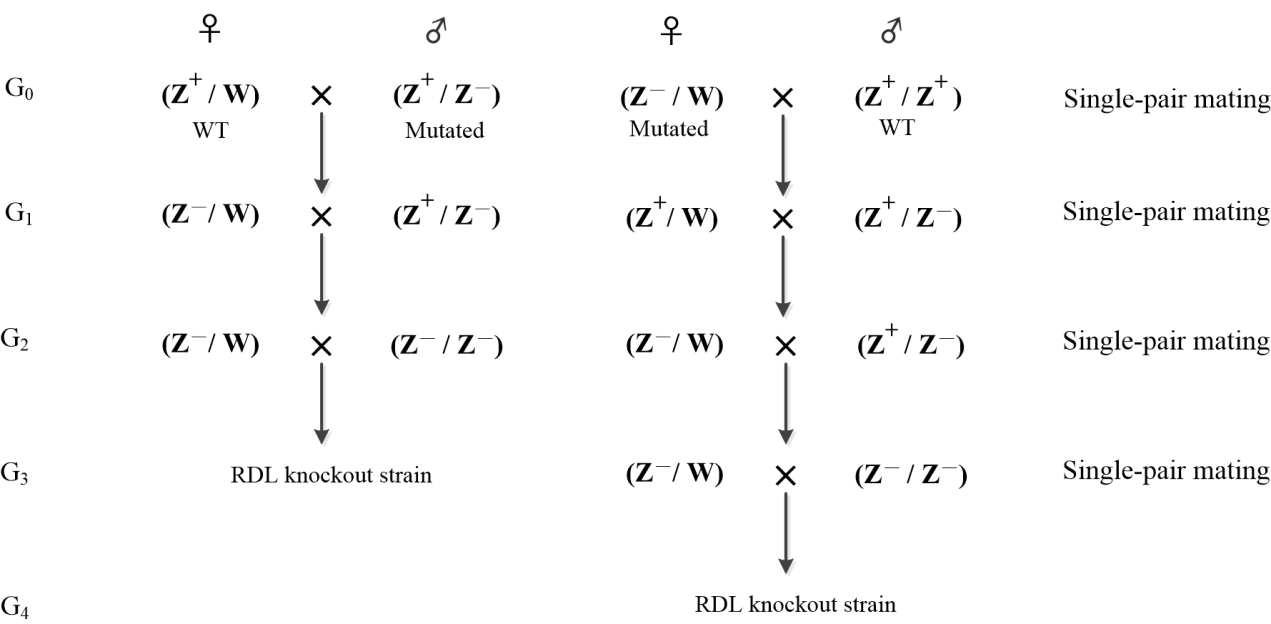
